# Distinct types of gut dysbiosis and oral–gut microbial signatures differentiate *C. difficile*, *Campylobacter*, and *Salmonella* infections: a cross-sectional study

**DOI:** 10.64898/2026.06.15.732290

**Authors:** Aleksander Mahnič, Rene Markovič, Marko Marhl, Andrej Golle, Nejc Stopnišek, Maja Rupnik

## Abstract

Gastrointestinal bacterial infections are associated with substantial perturbations of the gut microbiota, yet most microbiome studies have examined individual pathogens in isolation, limiting identification of shared and pathogen-specific microbial signatures across enteric infections. We performed a comparative analysis of fecal microbiota profiles from 586 stool samples using 16S rRNA gene sequencing, encompassing infections caused by *Clostridioides difficile*, *Campylobacter* spp., and *Salmonella* spp., alongside viral gastroenteritis, diagnostic-negative samples, and healthy controls. Fecal calprotectin concentrations were measured in a subset of samples to assess intestinal inflammation. Comparative analyses revealed two major dysbiotic configurations among bacterial enteric infections. *C. difficile* infection was characterized by an Enterococcus-dominated community structure, whereas *Campylobacter* and *Salmonella* infections were associated with enrichment of a tightly correlated consortium of oral-associated taxa, including *Streptococcus*, *Granulicatella*, and *Haemophilus*. These taxa were among the most informative features in an XGBoost machine-learning classifier, which accurately discriminated bacterial infection types from one another and from viral infections and healthy-associated microbiota profiles (macro F1 score = 0.74). In contrast, expansion of Enterobacteriaceae represented a shared, non-specific signature of intestinal disturbance and was associated with elevated fecal calprotectin concentrations. Together, these findings demonstrate that bacterial enteric infections are associated with distinct microbiota configurations that distinguish pathogen-specific signatures from general infection-related dysbiosis. The enrichment of oral-associated taxa in *Campylobacter* and *Salmonella* infections suggests a potential role for the oral–gut microbial axis in bacterial gastroenteritis and provides a foundation for future mechanistic studies and the development of microbiota-informed diagnostic, preventive, and therapeutic strategies.

**Author Summary:** Bacterial infections of the intestine can cause severe diarrhea and inflammation, but their effects on the community of microbes living in the gut are not fully understood. Most previous studies have focused on a single disease-causing organism, making it difficult to determine which microbiome changes are shared across infections and which are specific to particular pathogens. In this study, we compared gut microbiota profiles from people infected with *Clostridioides difficile*, *Campylobacter*, or *Salmonella*, and examined these alongside samples from individuals with viral gastroenteritis, diagnostic-negative diarrhea, and healthy controls. We found that bacterial enteric infections were associated with two distinct patterns of microbiome disruption. *C. difficile* infection was linked to an overgrowth of *Enterococcus*, whereas *Campylobacter* and *Salmonella* infections were characterized by increased levels of several bacterial groups that are commonly found in the mouth. In contrast, expansion of Enterobacteriaceae was observed across different infections and appeared to reflect general intestinal disturbance rather than a specific pathogen. Our findings identify microbial signatures that distinguish different bacterial infections and suggest that bacteria originating from the oral cavity may play an important role in some forms of gastroenteritis. These results provide a foundation for future studies aimed at understanding how microbial communities influence intestinal infection and recovery.

## INTRODUCTION

Gastrointestinal infections caused by bacterial pathogens represent a major global health burden, particularly in vulnerable populations. The gut microbiota plays a dual role in enteric infections, serving both as a target of pathogen-induced disruption and as a mediator of colonization resistance [1]. Dysbiosis in human gut has been consistently reported in infectious diarrhea [2], with *C. difficile* being well studied in this respect [3]. On the contrary, the understanding of such mechanisms in *Campylobacter* and *Salmonella* infections remains limited to small human cohort or clinical studies and studies including only children [4–7].

Development of *Clostridioides difficile* infection is typically associated with disturbed gut microbiota, characterized by depletion of obligate anaerobes and expansion of *Enterococcus* [3, 8]. In contrast, *Campylobacter* and *Salmonella* infections occur in immunocompetent individuals with eubiotic microbiota and are associated with acute mucosal inflammation and subsequent blooms of facultative anaerobes [4, 9]. Despite these mechanistic differences, infection-specific microbiota signatures have not been systematically compared.

Emerging evidence highlights an important role of the oral–gut axis, particularly the ectopic colonization of the gut by oral-associated bacteria such as *Streptococcus*, *Veillonella*, and *Haemophilus*, which may contribute to inflammatory gut perturbances, including in IBD and acute gastroenteritis [10–12]. However, although acute gastrointestinal infections can share inflammatory features with chronic inflammatory disorders (e.g., transient oxygenation and expansion of facultative anaerobes), the extent to which oral-to-gut translocation contributes to microbiota signatures during acute bacterial gastroenteritis remains insufficiently explored.

Our objective was to determine whether bacterial infections exhibit distinct microbiota profiles and to identify microbial markers that inform mechanistic understanding. We investigated infection-specific gut microbiota alterations in a collection of diagnostic residual samples with laboratory-confirmed *C. difficile*, *Campylobacter* spp., and *Salmonella* spp. infections. These were compared against an extensive set of reference groups, including viral infections (diagnostic samples with confirmed rotavirus and norovirus), diagnostic-negative samples, and healthy controls. We applied XGBoost machine learning with SHAP-based feature importance to assess the discriminative capacity of microbiota profiles and integrated correlation network analysis with differential abundance testing to define pathogen- and infection-specific signatures.

## METHODS

### Sample collection

A total of 539 residual diagnostic stool samples were included in this study. Samples were distributed into different study groups based on routine diagnostic testing. Samples were obtained at three sites of the National Laboratory of Health, Environment and Food (NLZOH): Maribor, Celje, and Kranj, during the period August 1st 2023 to January 31st 2025. The test groups included samples tested positive for one of three target bacterial gastrointestinal pathogens: *Clostridioides difficile*, *Campylobacter* spp., and *Salmonella* spp.. Samples that tested positive for either rotavirus or norovirus (viral infections) or negative for all gastrointestinal pathogens examined (negative controls) were included as comparative study groups. Moreover, healthy controls (*n* = 47) were obtained from individuals residing in the same geographic region and collected as part of a previously published study (Mahnic et al., 2018). The only inclusion criterion was age of 4 years or more. Due to ethical and logistical constraints, only age and gender were recorded as metadata. The study was approved by the institutional ethical committee (Medical Faculty, University of Maribor; 038/2023/5-513). Informed consent was not obtained, as the study was conducted on residual diagnostic samples collected in the laboratory, with no direct contact with patients.

### Pathogen detection

Pathogen detection was performed at three regional NLHEF laboratories, each applying ISO-standardized diagnostic workflows adapted to locally available instrumentation. Across all sites, *Clostridioides difficile* was detected by selective cultivation on CHROMID® *C. difficile* agar (bioMérieux, Marcy-l’Étoile, France), followed by identification of presumptive colonies using matrix-assisted laser desorption/ionization time-of-flight mass spectrometry (MALDI-TOF MS; Bruker Daltonics, Billerica, MA, USA). Detection of *C. difficile* toxin genes was performed using the Alethia® *C. difficile* (Meridian Bioscience, Cincinnati, OH, USA) directly from stool samples. In cases where stool testing was negative, but culture yielded *C. difficile*, toxin gene detection was additionally performed on cultured isolates.

*Salmonella* spp. and *Campylobacter* spp. were detected by cultivation on IRIS® Salmonella Agar (bioMérieux, Marcy-l’Étoile, France) and Campylobacter Agar Base Blood-Free (modified charcoal cefoperazone deoxycholate agar [mCCDA], Oxoid, Thermo Fisher Scientific, Basingstoke, UK), respectively, following ISO guidelines. Presumptive isolates were identified to the species level using MALDI-TOF MS (Bruker Daltonics, Billerica, MA, USA).

Viral pathogens, including norovirus (genogroups I and II) and rotavirus, were detected by real-time reverse transcription PCR (RT-PCR) using LightMix® Modular assays (TIB Molbiol GmbH, Berlin, Germany). Nucleic acids were extracted from stool samples using commercially available viral nucleic acid extraction kits, according to the manufacturers’ instructions, and amplified on compatible real-time PCR platforms.

### 16S rRNA gene amplicon sequencing

Total DNA was extracted from stool samples using the PowerFecal Kit (QIAGEN) according to the manufacturer’s recommendations. The V3–V4 hypervariable region of the bacterial 16S rRNA gene was amplified using primers Bakt_341F (5’-CCTACGGGNGGCWGCAG-3’) and Bakt_805R (5’-GACTACHVGGGTATCTAATCC-3’). Sequencing was performed on the Illumina NextSeq platform (2 × 300 bp). Raw reads were quality filtered and processed into amplicon sequence variants (ASVs) using mothur (v1.48.0). Taxonomic assignment was carried out using the RDP reference database (trainset v.9). Negative sequencing controls included extraction, library preparation, and sequencing blanks. Positive controls consisted of an in-house 24-member mock bacterial community representing typical gut microbiota diversity. Sequencing data are available in the NCBI Sequence Read Archive under accession PRJNA1469393 and MG-RAST project access number mgp85661 (Heatlhy controls) [13].

### Fecal calprotectin quantification

Fecal calprotectin was quantified directly from stool using approximately 50 µL of sample. Measurements were performed using the CalprestTurbo latex turbidimetric assay (Eurospital Diagnostic) following the manufacturer’s instructions. Analyses were run on the Roche Cobas c111 turbidimetric analyzer. The lower limit of detection was 20 mg/kg, as specified by the manufacturer.

### Data preparation

ASV count data and sample metadata were loaded and inner-joined on sample identifiers. Prior to model training, ASV count data were transformed using the centered log-ratio (CLR) transformation to account for the compositional nature of microbiome data. A pseudo count of 0.1 was added before transformation to handle zero values. Features consisted exclusively of CLR-transformed ASV columns, and the target variable was read from a designated metadata column and integer-encoded.

To address class imbalance, random oversampling was applied exclusively to the training partition within each cross-validation fold, ensuring that the validation set always reflected the natural class distribution.

An XGBoost gradient-boosted tree classifier was trained and evaluated using stratified 5-fold cross-validation. Per-fold predictions, class probabilities, and feature importances were collected; aggregate metrics — overall accuracy, per-class precision, recall, F1-score, and a pooled confusion matrix — were computed across all folds. Additionally, per-class and macro-averaged AUROC and AUPRC were computed for each fold using one-vs-rest binarization and averaged across folds. Hyperparameters were determined by prior grid search and set as follows for the final analysis: n_estimators = 500; max_depth = 4; learning_rate = 0.1; subsample = 0.9; colsample_bytree = 0.9.

Per-fold SHAP (SHapley Additive exPlanations) values were computed on held-out validation samples using the unified SHAP Explainer API, yielding three-dimensional arrays (samples × features × classes). Feature importance was summarized as the mean absolute SHAP value averaged across samples, classes, and folds, and reported for the top 20 features. For exploratory visualization, t-SNE (perplexity = 30, PCA initialization) was applied to standardized ASV features across all samples, and cross-validation predictions were overlaid on the embedding to illustrate classification performance in low-dimensional space.

The final model was trained on the complete dataset with oversampling applied, using the same hyperparameters, and saved for downstream inference.

### Statistical analysis

All statistical analyses, machine-learning procedures, and visualizations were conducted in R (R Foundation for Statistical Computing). All analysis scripts are available at GitHub repository https://github.com/ReneMarkovic/ASV-Genie.

Alpha diversity was calculated using the Shannon index. Beta diversity was assessed using ANOSIM with Bray–Curtis dissimilarities implemented in the *vegan* package. Pairwise comparisons were performed using the Wilcoxon rank-sum test, and multiple-group comparisons using the Kruskal–Wallis test. Correlation networks were inferred using the NetCoMi package[14], applying spring-based correlation metrics. Differentially abundant taxa were identified using ALDEx2. Where required, Benjamini–Hochberg false discovery rate (FDR) correction was applied, with FDR-adjusted p < 0.05 considered significant. Partitioning around medoids (PAM, R package ‘cluster) approach was implemented using Bray-Curtis dissimilarities to distinguish community types beyond our available metadata.

For machine learning, both Random Forest and XGBoost classifiers were evaluated; XGBoost was selected based on superior performance (macro F1 score 0.49 and 0.53 for Random Forest and XGBoost, respectively). ASVs corresponding to the target bacterial pathogens (*C. difficile*, *Salmonella*, and *Campylobacter*) were removed from the model input to focus on predictive contributions of the remaining microbiota. Oversampling was applied during test-set creation to address class imbalance. Model performance was assessed using 5-fold cross-validation. The XGBoost model utilized the optimized hyperparameter profile established during the initial grid search. Feature importance for each target class was determined using SHAP (SHapley Additive exPlanations) values.

## RESULTS

Bacterial infection–specific gut microbiota characteristics were investigated using stool samples submitted for routine diagnostic testing for gastrointestinal infection. To contextualize bacterial infection–associated shifts, microbiota profiles from samples positive for bacterial pathogens were compared with (i) samples with confirmed rotavirus or norovirus infection (viral infections), (ii) diagnostic-negative samples in which no pathogen was detected (negative controls) and (iii) healthy volunteers from general population living in the same geographic area (healthy controls). Notably, diagnostic-negative samples were obtained from symptomatic individuals for whom pathogen testing was clinically indicated; thus, unmeasured contributors (e.g., non-tested pathogens, non-infectious etiologies, comorbidities, or medication use) may be present.

Analysis focused on three bacterial pathogens: *Clostridioides difficile*, *Campylobacter* spp., and *Salmonella* spp., and compared their microbiota profiles with healthy controls, diagnostic negative samples, and viral infection samples (Table 1). The median participant age was 55 years; participants with *C. difficile* infection were older (median 69 years; Kruskal–Wallis test, *p* < 0.001), whereas no age differences were observed among the remaining infection groups. Gender distribution varied across groups but did not reach statistical significance (Fisher’s exact test, *p* > 0.05 for all comparisons).

**Table 1.**
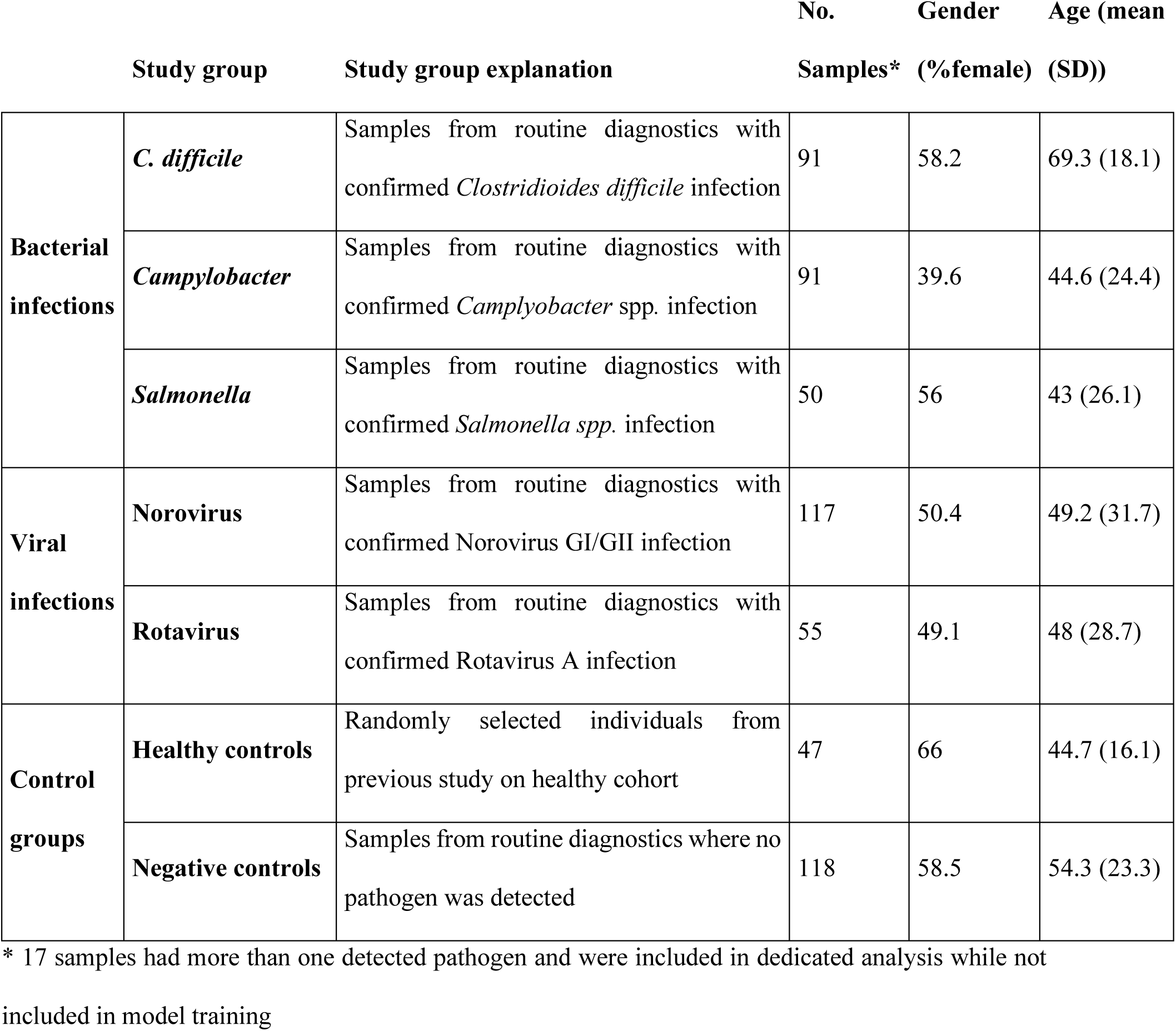
Cohort characteristic according to study group.

### Global microbiota structure and community types

To enable interpretation of bacterial infection-specific signatures in the context of broader community variation, global community structure was assessed. Samples span from healthy-associated profiles to two dysbiosis-like compositions (Fig. 1A; healthy controls highlighted in dark blue). Partitioning around medoids (PAM) supported a three-cluster solution (community types, CT1–CT3; silhouette index = 0.123; Fig. 1A).

**Fig. 1.**
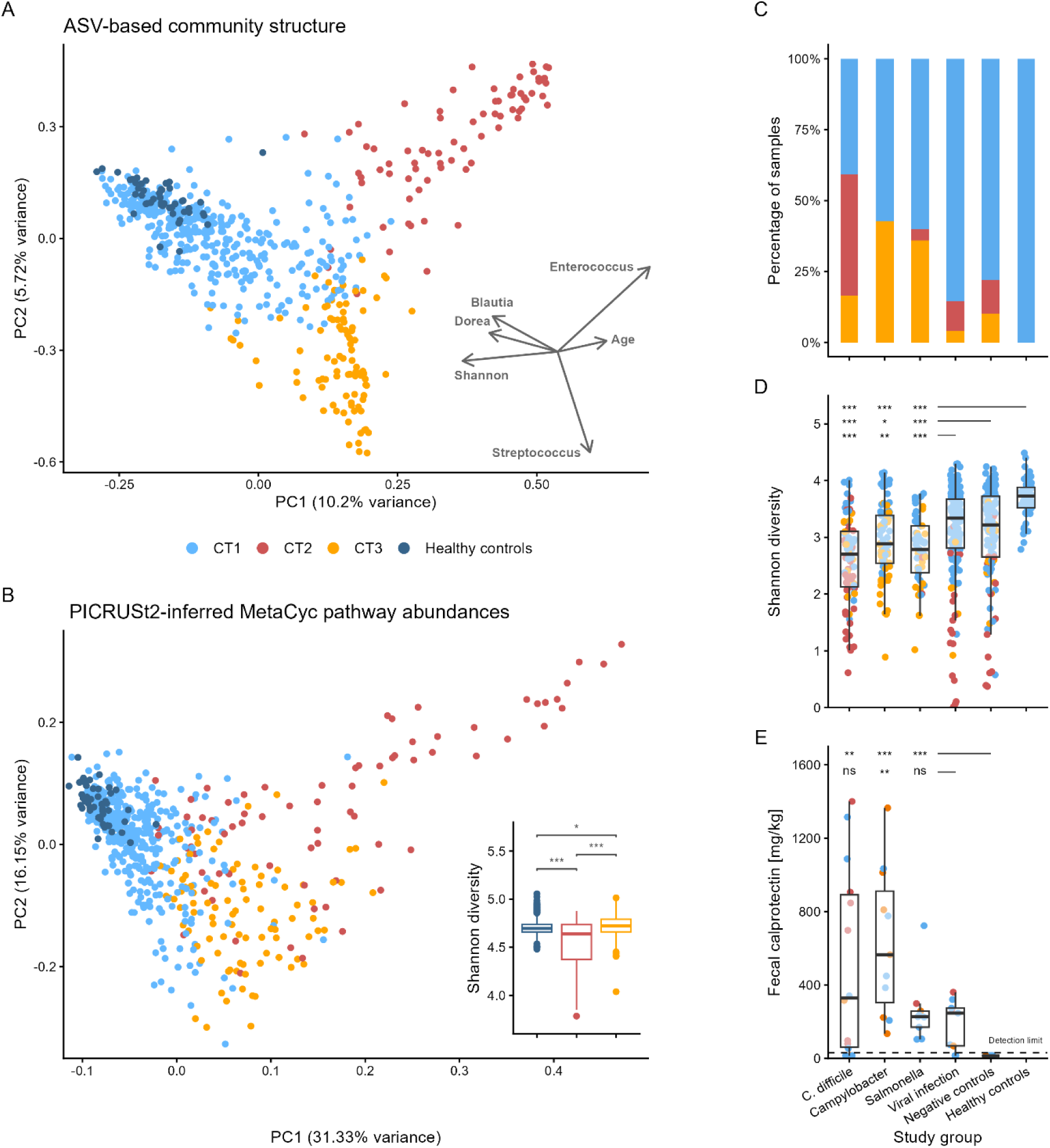
Gut microbiota characteristics and fecal calprotectin in different bacterial infections. (A) PCA visualization based on ASV-level community structure, color-coded by community type designation, with healthy controls additionally highlighted in dark blue (CT1). Arrows present direction of significant (*p* < 0.05) bacterial community and metadata factors which display *R*^2^ > 0.25. (B) PCA visualization based on predicted KEGG pathways, color-coded by community type designation, with healthy controls additionally highlighted in dark blue (CT1). Nested plot presents Shannon diversity calculated from predicted KEGG pathways. (C) Distribution of study groups according to community type designation. (D) Bacterial community diversity (Shannon index) according to study group. (E) Fecal calprotectin levels according to study group. Significance is denoted by following legend: > 0.05: ns, 0.05 – 0.01: *, 0.01 – 0.001: **, < 0.001: ***.

Across all infection-associated study groups, *Enterobacteriaceae* were consistently elevated compared with healthy controls (*p* < 0.05 for all comparisons), while other differentially abundant taxa showed cluster-specific associations. CT1 was the dominant community type (*n* = 417, 71.1%) and most closely resembled healthy microbiota, although *Enterobacteriaceae* were still increased relative to healthy controls. CT2 (*n* = 76, 13.0%) was characterized by increased *Enterococcus* abundance. CT3 (*n* = 93, 15.9%) showed increased abundance of oral-associated genera, most prominently *Streptococcus* ASVs (Supplementary Fig. S1). CT2 was associated with older age (Kruskal–Wallis test, *p* < 0.001), whereas age did not differ between CT1 and CT3 (Wilcoxon test, *p* = 0.538).

Community types only partially overlapped with infection types (Fig. 1C). *C. difficile* infection was enriched in CT2 compared with other groups (42.9% vs. 6.4%; Fisher’s exact test, *p* < 0.001). In contrast, *Campylobacter* and *Salmonella* infections were infrequent in CT2 and were more likely to cluster in CT3 compared with other diagnostic groups (42.6%, 36.0%, and 8.4% for *Campylobacter*, *Salmonella*, and all other study groups, respectively; Fisher’s exact test, *p* < 0.001 for all comparisons). All three bacterial infection groups exhibited reduced alpha diversity relative to both control groups (negative and healthy) and viral infections (Shannon index; Wilcoxon test, *p* < 0.02 for all comparisons; Fig. 1D). Furthermore, within bacterial infections, samples assigned to CT2 or CT3 had lower diversity than samples with the same infection assigned to CT1 (Wilcoxon test, *p* < 0.001). Cluster CT1 comprised samples from all infection and control groups, indicating that presence of infection was not necessarily also associated with global microbiota alterations.

Community types also differed in inferred metabolic potential (PICRUSt2). Changes in inferred metabolic potential to a large extent mirrored those observed based on community structure, indicating not only taxonomical shift, but also alterations in functional potential of these communities (Fig. 1B). The change in predicted functions was not confined to a single pathway class but spanned multiple metabolic modules. The most strongly shifted PICRUSt2-inferred MetaCyc pathway abundances showed a progressive deviation from healthy-toward CT2/CT3-associated disruption. These shifts included depletion of pathways typically associated with anaerobic, community-level metabolism (e.g., Wood–Ljungdahl pathway, vitamin B12 salvage, and acetoclastic methanogenesis) alongside enrichment of pathways consistent with increased biosynthetic autonomy (e.g., de novo purine and pyrimidine nucleotide biosynthesis, L-lysine biosynthesis, and NADPH-generating routes) (Supplementary Fig. S2).

Fecal calprotectin was used as a surrogate marker of intestinal inflammation (Fig. 1E). Calprotectin levels were higher in all bacterial and viral infections than in diagnostic-negative controls (Wilcoxon test, p < 0.005). *Campylobacter*-positive samples elicited a significantly higher calprotectin response (mean 632 mg/kg) than *Salmonella*-positive samples (mean 260 mg/kg; Wilcoxon test, *p* = 0.038). Meanwhile, intermediate calprotectin levels were observed in *C. difficile*-positive samples (mean 513 mg/kg), which showed no statistically significant deviation from either group (Kruskal–Wallis test, *p* = 0.255).

### Predictive microbiota signatures of infection

Since global dysbiosis patterns only partially corresponded to infection status and may reflect unmeasured factors (e.g., antibiotic exposure or gastrointestinal comorbidities), machine learning was used to identify infection-associated microbiota features beyond global community shifts. An XGBoost classifier was trained to discriminate among study groups using ASV profiles. Overall predictive performance was modest (macro F1 score = 0.49). Performance did not differ across community types (accuracy 46.7%, 52.1%, and 45.1% for CT1, CT2, and CT3, respectively; Fisher’s exact test, p > 0.05 for all comparisons).

Performance, however, differed markedly across study groups (Fig. 2A–B). *C. difficile* infection showed the most distinct microbiota profile (F1 score = 0.68), whereas *Campylobacter* and *Salmonella* were predicted with notably lower accuracy (F1 score = 0.46 and 0.30, respectively). For *Campylobacter*, pathogen relative abundance was weakly positively correlated with model-predicted infection probability (Spearman’s rho = 0.210, p = 0.046; Supplementary Fig. S3), indicating pathogen-load–associated shifts in community disruption. This association was not observed for *Salmonella* or *C. difficile* infection. Other than bacterial infections, healthy controls were predicted with high accuracy (F1 score = 0.76), while diagnostic negatives and viral infection samples showed lower performance (F1 score = 0.35, 0.44, and 0.45 for diagnostic negatives, rotavirus, and norovirus, respectively; Fig. 2B). Misclassifications occurred most frequently between diagnostic negatives and viral infections, and between *Campylobacter* and *Salmonella* infections (Fig. 2D), consistent with partially overlapping microbiota signatures (Fig. 2A).

**Fig. 2.**
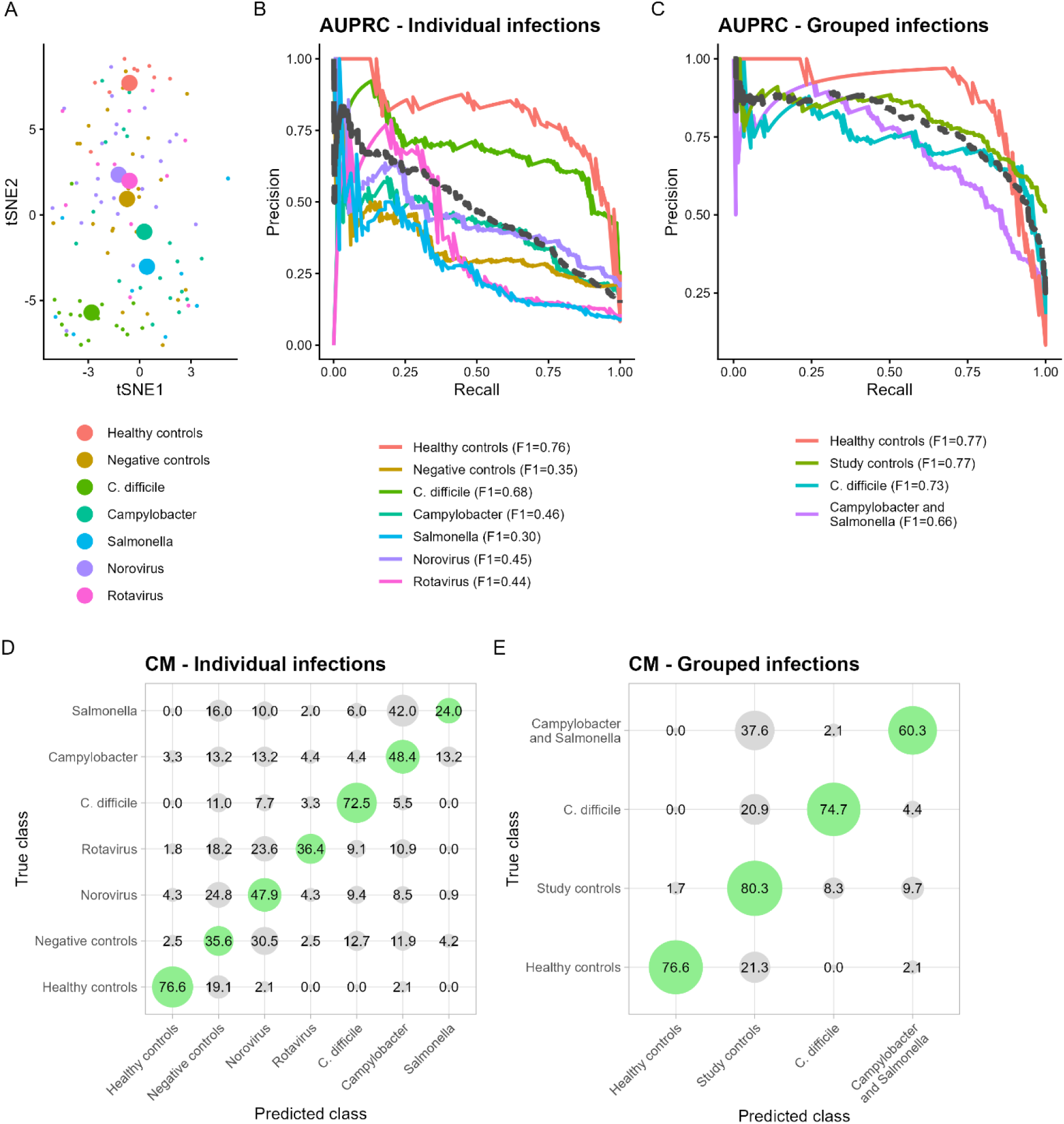
Model prediction of infection groups based on gut bacterial community structure. Models were built implementing XGBoost approach with 5-fold validation. (A) t-SNE visualization based on XGBoost predictions. (B) AUPRC curves of XGBoost model color coded by study group. Black dotted line presents model average. (C) AUPRC curves of XGBoost model color coded by study groups after study groups were merged, highlighting improved overall model power after merger. More specifically, study controls include negative samples and viral infections and another merged group presents combined *Campylobacter* and *Salmonella* infections. (D) Confusion matrix obtained from XGBoost model for individual study groups (E) Confusion matrix obtained from XGBoost model for grouped infections.

To test whether misclassification reflected shared community-level features, an additional model was trained after combining (i) *Campylobacter* and *Salmonella* into a single group and (ii) diagnostic negatives and viral infection samples into a second combined group. This model showed substantially improved overall performance (macro F1 score = 0.74), with all target groups exceeding an F1 score of 0.65 (Fig. 2C). In this reduced-label model, bacterial infections (i.e., *C. difficile* and combined *Campylobacter*/*Salmonella*) and healthy controls were distinguished with high accuracy, whereas most remaining misclassifications involved the combined diagnostic negatives and viral infections group (Fig. 2E).

### Samples with more than one detected pathogen

During sampling we obtained seventeen specimens, where two or more of the five target pathogens were detected. Most of these involved concomitant *C. difficile* and a viral infection (*n* = 10), while other combinations were infrequent (Supplementary Table S1). Samples with more than one detected pathogen were excluded from model training and the trained classifier was applied post hoc to these samples.

Among samples that included *C. difficile* and other pathogens, the model predicted *C. difficile* in 66.7% (8/12) of samples, increasing to 70% (7/10) when the concomitant detected pathogen was viral. In samples with concomitant bacterial and viral pathogen, the model predicted the viral infection in 3/11 instances, suggesting that viral infection contributed less frequently to the dominant microbiota signature in these cases. Seven samples included *Campylobacter* or *Salmonella* together with another pathogen, but the model did not predict either pathogen in any of these samples. Although the sample size is too small for robust inference, this pattern suggests that *Campylobacter*/*Salmonella*-associated microbiota signatures may be masked when co-occurring with other strong dysbiosis drivers (Supplementary Table S1).

### Bacterial infection–specific microbiota characteristics

To identify robust bacterial infection–associated features, microbiota profiles from *C. difficile* positive samples (*n* = 91) and combined *Campylobacter*/*Salmonella* positive samples (*n* = 141) were compared against (i) healthy controls (*n* = 47), (ii) diagnostic-negative samples (n = 118), and (iii) viral infections (rotavirus and norovirus combined; *n* = 172) using population level analysis.

Most retained ASVs formed a large health-associated cluster (Fig. 3, yellow nodes) comprising taxa depleted across all bacterial infections, with the strongest depletion observed in *C. difficile* samples. These included primarily different representatives from *Blautia, Lachnospiraceae, Bifidobacterium, Clostridium* XIVa and other health-associated *Bacillota.* In addition, a distinct cluster (Fig. 3, blue nodes) was enriched for oral-associated taxa and increased in *Campylobacter*/*Salmonella* infections. Infection-associated ASVs identified from the network were concordant with two complementary feature-selection approaches: (i) SHAP feature ranking from the XGBoost classifier and (ii) ALDEx2 differential abundance testing.

**Fig. 3.**
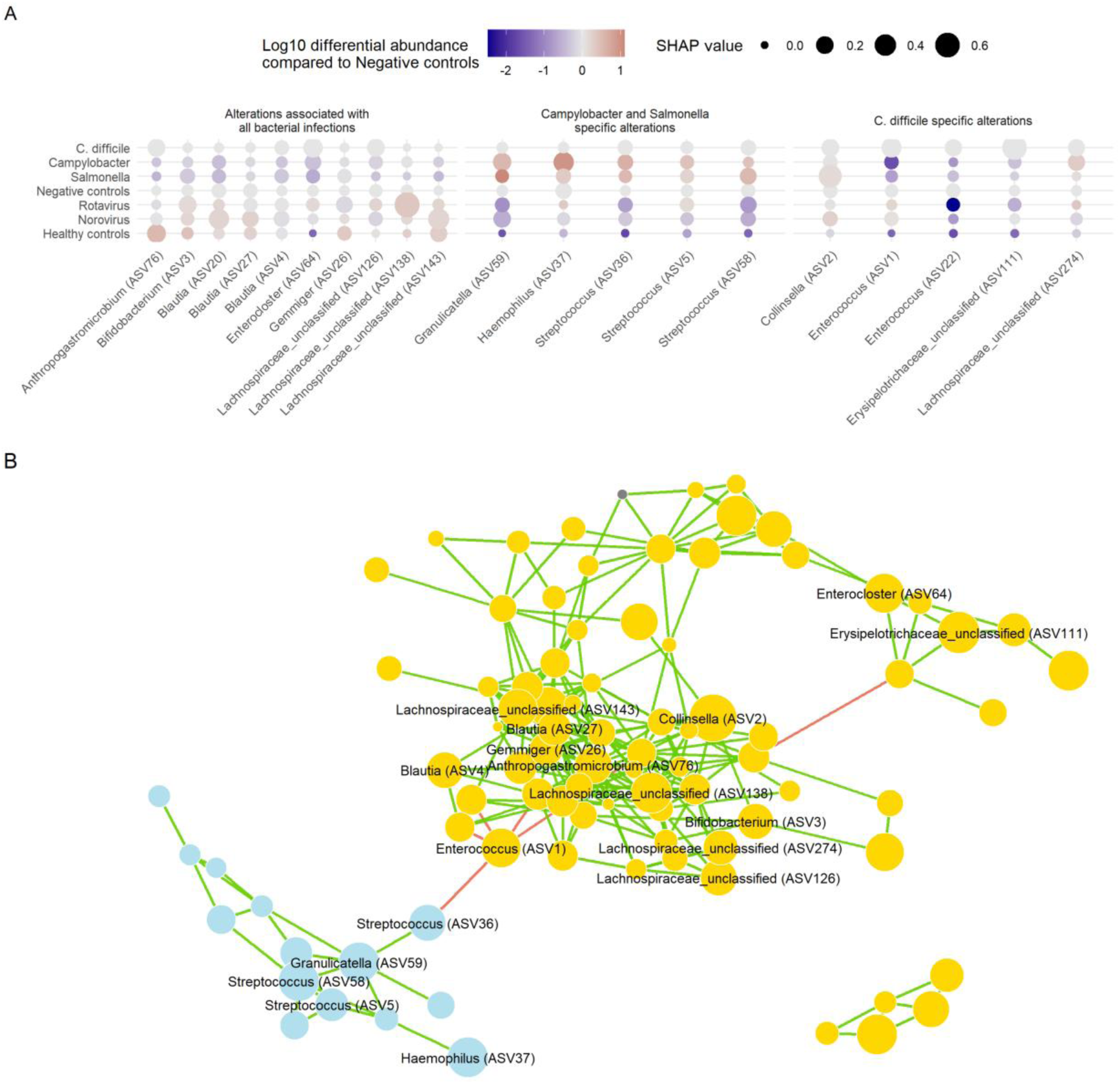
Bacterial infection–specific alterations in gut microbiota composition. (A) The highlighted ASVs represent features consistently identified as significant by both the machine learning approach (XGBoost with SHAP-based feature ranking) and the differential abundance analysis (ALDEx2). From these, the taxa shown are those ranked among the top 30 features with SHAP and simultaneously exhibiting significant differential abundance in either the *C. difficile* group or the combined *Campylobacter/Salmonella* group, compared with both control groups and viral infections (ALDEx2, FDR < 0.05). For each ASV, the figure displays the study-group–associated SHAP value (circle size) and the log10 fold change relative to healthy controls (color gradient, red denoting increase in infection and blue decrease in infection as compared to negative samples). (B) Taxa enriched *Campylobacter/Salmonella*–associated communities are positively correlated with each other, as indicated by the correlation network analysis. Cluster associated with *Campylobacter/Salmonella* infection is highlighted with blue color of nodes while others are presented in yellow. Additionally, size of nodes represents SHAP value of individual ASVs included in the correlation network.

### *Campylobacter*/*Salmonella*–specific microbiota alterations

Sixteen ASVs were increased in *Campylobacter*/*Salmonella* infections compared with both control groups and viral infections combined (ALDEx2, we.eBH < 0.05), corresponding to an average 14.2-fold increase (SD = 2.1). Ten of these ASVs clustered in the correlation network, forming a distinct module (Fig. 3, blue nodes). This module was enriched for taxa classified as *Streptococcus* (n = 9), *Haemophilus*, *Granulicatella*, *Veillonella*, *Rothia*, *Peptostreptococcus*, and *Lactobacillus*. ASVs within this module were strongly positively correlated, consistent with a co-occurring community signature and/or increased representation of oral-associated taxa in stool.

Five ASVs from this module ranked among the top 30 model features (three *Streptococcus* ASVs, *Haemophilus*, and *Granulicatella*). Group-specific SHAP values indicated that *Haemophilus* (ASV37) and *Granulicatella* (ASV59) contributed most to *Campylobacter* prediction, whereas *Streptococcus* ASV58 was the strongest predictor for *Salmonella* infection (Fig. 3).

The cumulative relative abundance of the oral-associated module was higher in most *Campylobacter*/*Salmonella* samples with consequential profound effect on predicted metabolic potential of these communities. Metabolic prediction linked oral genera to a denitrification–heme–fermentation–NAD signature, indicating repurposing of oral biofilm pathways (nitrate respiration, heme-dependent respiration, mixed-acid/2,3-butanediol fermentation) for survival in the inflamed gut, thereby exploiting iNOS-derived nitrate and dysbiotic substrates during *Campylobacter*/*Salmonella* infections.

Five ASVs from this cluster also ranked among the top 30 model features (three *Streptococcus* ASVs, *Haemophilus*, and *Granulicatella*). Group-specific SHAP values indicated that *Haemophilus* (ASV37) and *Granulicatella* (ASV59) contributed most to *Campylobacter* prediction, whereas *Streptococcus* ASV58 was the strongest predictor for *Salmonella* infection (Fig. 3).

### *C. difficile*–specific microbiota alterations

*C. difficile*–associated changes were not concentrated in a single correlated module to the same extent as the *Campylobacter*/*Salmonella* signature. Increases were observed for *Enterococcus* (ASV1 and ASV22), *Erysipelotrichaceae* (ASV111), and *Clostridium* XIVa group (ASV64), alongside additional enrichment of *Actinobacteriota*- and *Bacillota*-affiliated taxa (including *Eggerthella*, Erysipelotrichaceae, and Lachnospiraceae), although not all exceeded multiple-testing correction.

A strong co-occurrence between *C. difficile* positivity and *Enterococcus* was observed: 91.2% of *C. difficile* samples contained *Enterococcus* versus 42.6% of non–*C. difficile* samples (Fisher’s exact test, p < 0.001). *Enterococcus* ASVs (ASV1 and ASV22) were more abundant in *C. difficile* infections and were most prominent in samples assigned to community type CT2. *Enterococcus* ASV1 abundance increased with age (Pearson r = 0.341, p < 0.001), whereas the *C. difficile*–associated ASV62 did not correlate with age (r = −0.002, p = 0.983) and did not correlate with *Enterococcus* abundance (r = −0.040, p = 0.738). In contrast, *C. difficile* relative abundance correlated positively with *Erysipelotrichaceae* ASV111 (Pearson r = 0.161, p < 0.001), with this ASV showing also higher prevalence in *C. difficile* positive samples (71.4% vs. 45.6%; Fisher’s exact test, p < 0.001). Consistent with these results, SHAP ranking identified *Erysipelotrichaceae* ASV111 as the strongest positive predictor of the *C. difficile*–associated microbiota disruption.

## Discussion

The gut microbiota plays a pivotal role in the pathogenesis of gastrointestinal infections and represents a promising target for both therapeutic and preventive strategies [15, 16]. However, identification of infection-specific microbiota signatures is often restricted to comparisons between a single infection and healthy controls, which substantially limits the power and robustness of the resulting findings [17]. In this study, we aimed to identify infection-specific microbiota signatures across three bacterial pathogens: *C. difficile*, *Campylobacter* spp., and *Salmonella* spp. While there is substantial evidence describing gut microbiota alterations associated with *C. difficile* infection [3, 18, 19], data for the other two bacterial infections remain comparatively limited. To address this gap, we included multiple bacterial infections alongside a comprehensive set of reference groups (viral infections, diagnostic negatives, and healthy controls). Coupled with machine learning approaches, this design enabled a more robust and comparative investigation of infection-specific microbiota signatures.

Broad analysis demonstrated the commonly reported dysbiosis-associated increase in *Enterobacteriaceae* across all study groups in our cohort, including diagnostic negatives, relative to healthy controls. This shift therefore reflects a general disruption rather than a disease-specific alteration. Expansion of *Enterobacteriaceae* is a well-established consequence of inflammation, associated with loss of colonization resistance, generation of reactive oxygen and nitrogen species that can serve as terminal electron acceptors, and increased luminal oxygen availability, collectively favoring facultative anaerobes over strictly anaerobic commensals from *Bacillota* and *Bacteroidota* [1, 20–22].

Furthermore, our findings suggest that gut microbiota disruption follows two distinct patterns, with different gastrointestinal pathogens showing preferential, though not exclusive, associations with these configurations. As previously proposed, distinct dysbiotic states are linked to specific physiological alterations that shape disease susceptibility and progression [17, 23]. In our cohort, microbiota composition transitioned from a healthy-associated state (CT1) to either an *Enterococcus*-dominated configuration (CT2) or a state characterized by increased abundance of a tightly correlated consortium of oral-associated taxa (CT3). Both CT2 and CT3 exhibited reduced community diversity relative to CT1. Similar to microbiota composition, the most strongly altered PICRUSt2-inferred MetaCyc pathway abundances also demonstrated a progressive deviation from the healthy-associated state toward CT2/CT3-linked disruption. These changes were characterized by depletion of pathways typically associated with anaerobic, community-level metabolism (e.g., the Wood–Ljungdahl pathway, vitamin B12 salvage, and acetoclastic methanogenesis), alongside enrichment of pathways indicative of increased biosynthetic autonomy (e.g., de novo purine and pyrimidine nucleotide biosynthesis, L-lysine biosynthesis, and NADPH-generating pathways). Collectively, these inferred functional shifts are consistent with a transition from a microbiota with greater reliance on cross-feeding and metabolite exchange toward one exhibiting increased biosynthetic self-sufficiency [24]. However, this interpretation should be considered with caution, as it is based on predicted metabolic potential rather than direct functional measurements.

Notably, fecal calprotectin levels did not differ between community types but were instead more strongly associated with the pathogen, independent of community structure. Mean calprotectin levels in bacterial infections were consistent with clinically relevant intestinal inflammation according to British Society of Gastroenterology guidance and were significantly elevated compared to diagnostic negative controls across all targets bacterial infections. Pathogen-specific associations were also evident: *C. difficile* was significantly linked to CT2, whereas *Campylobacter* and *Salmonella* infections were predominantly associated with CT3.

In contrast to *Salmonella* and *Campylobacter* gastroenteritis, which can arise in individuals without disrupted gut microbiota, *Clostridioides difficile* infection (CDI) typically occurs in the context of a pre-disturbed microbiota [25]. CDI risk is strongly associated with recent broad-spectrum antibiotic use, which disrupts colonization resistance and facilitates *C. difficile* overgrowth [26] and could therefore be an important confounding factor leading to unique microbiota signatures observed for this pathogen. Antibiotic-associated dysbiosis also creates ecological opportunities for expansion of multidrug-resistant enterococci. Importantly, the relationship between *C. difficile* and enterococci extends beyond simple niche co-occupation; current evidence indicates that enterococci can enhance *C. difficile* fitness and virulence through mechanisms such as mutual biofilm formation and metabolic cross-feeding [8, 27]. Consistent with this, a strong positive correlation between *C. difficile* and *Enterococcus* was observed in our cohort. However, an even stronger positive correlation was detected with a representative of the *Erysipelotrichaceae*, based on ASV sequence inspection most likely *Clostridium innocuum*, although further validation is required to confirm this assignment. This organism, while less frequently reported, is an established vancomycin-resistant opportunistic pathogen with demonstrated pathogenic potential [28, 29] and has previously been identified in patients with CDI and inflammatory bowel disease [30, 31]. In our dataset, while *Enterococcus* co-occurred with *C. difficile*, the *Erysipelotrichaceae* representative additionally exhibited a particularly strong positive correlation with *C. difficile* abundance.

Microbiota perturbations observed in *Salmonella* and *Campylobacter* infections, in contrast to CDI, largely reflect the infection process itself rather than prior antimicrobial exposure. Nonetheless, prospective evidence suggests that baseline microbiota composition may influence susceptibility to *Campylobacter* infection. In a human challenge study, individuals who became *Campylobacter*-positive exhibited distinct baseline fecal microbiota compared to those who remained negative, most notably with increased *Escherichia* and *Streptococcus* [6]. Similarly, a study on travelers at high risk for *Campylobacter* acquisition reported that individuals who later developed infection had lower pre-travel microbiota diversity [32]. Despite these findings, both *Campylobacter* and *Salmonella* are generally considered capable of causing infection in otherwise eubiotic microbiota, with infection risk driven primarily by exposure dose [33]. These two pathogens share key biological and ecological characteristics, including induction of diarrhea, mucosal inflammation, luminal oxygenation, and expansion of facultative anaerobes [34, 35]. Beyond this common expansion, our study identified a previously unreported feature: a significant increase in oral-associated taxa during infection with both pathogens. These included multiple *Streptococcus* species, *Granulicatella*, and *Haemophilus*, which also displayed strong positive correlations across individuals. Emerging evidence indicates that conditions involving mucosal inflammation and increased intestinal transit, most extensively studied in inflammatory bowel disease, are associated with enrichment of oral-derived taxa such as *Streptococcus*, *Veillonella*, *Haemophilus*, and *Granulicatella* [36, 37]. This phenomenon may be linked, although not exclusively, to oral health status, including plaque accumulation and subsequent periodontitis [38, 39]. Oral-to-gut bacterial translocation occurs continuously in healthy individuals but may be amplified during gastrointestinal disease due to altered luminal conditions, compromised mucosal barriers, or selective pressures favoring facultative anaerobes capable of thriving in inflamed or oxygenated environments [23, 38, 39]. Whether this represents a general consequence of infectious diarrhea or exhibits pathogen-specific patterns has not been previously explored in a multi-pathogen comparative framework.

To further elucidate the mechanisms underlying oral–gut translocation observed in our study, additional metadata including oral health status, comorbidities, and medication use, would be essential for host factor–resolved analyses. The absence of such data represents an important limitation, albeit one inherent to the study design, which only enabled large-scale sample collection without extended patient-level metadata. Interpretation is further constrained by the fact that these observations are based solely on stool samples. Definitive confirmation of oral–gut translocation would require strain-resolved analyses of paired oral and stool samples using metagenomic or culturomic approaches.

This study provides a comparative analysis of gut microbiota alterations across *C. difficile*, *Campylobacter*, and *Salmonella* infections, identifying both shared and pathogen-associated features of dysbiosis. While reported *C. difficile* association with *Enterococcus* largely consolidates previous findings, the enrichment of oral-associated taxa in *Campylobacter* and *Salmonella* infections reveals a previously underexplored aspect of infection-associated dysbiosis and supports a role for enhanced oral–gut microbial translocation. These insights have translational relevance, supporting the development of community-level microbiota-based diagnostics. Furthermore, the identification of oral microbiota as a potential upstream reservoir suggests novel preventative and therapeutic strategies, where targeting oral–gut microbial dynamics may reduce infection risk or alleviate disease severity.

## Acknowledgements

We would like to acknowledge the contributions of Živa Petrovič, Mateja Ravnik, and Tanja Blatnik from the National Laboratory of Health, Environment and Food (Center for Medical Microbiology) for their assistance with the organization of residual samples and collection of metadata.

## Data availability

Sequencing data are available in the NCBI Sequence Read Archive under accession PRJNA1469393 and MG-RAST project access number mgp85661 (Heatlhy controls).

